# *OnCorr:* A pan-cancer mRNA-protein correlation tool for precision oncology

**DOI:** 10.1101/2025.06.12.659426

**Authors:** Urwah Nawaz, Niantao Deng, Ori Livson, Chelsea Mayoh, Loretta M S Lau, Roger R Reddel, Bhavna Padhye, Rebecca C Poulos

**Affiliations:** ProCan®, Children’s Medical Research Institute, Faculty of Medicine and Health, The University of Sydney, Westmead, NSW, Australia; Children’s Cancer Institute, Lowy Cancer Research Centre, UNSW Sydney, Sydney, NSW, Australia; School of Clinical Medicine, UNSW Medicine & Health, UNSW Sydney, Sydney, NSW, Australia; Kids Cancer Centre, Sydney Children’s Hospital, Sydney, NSW, Australia; Cancer Centre for Children, The Children’s Hospital at Westmead, Westmead, NSW, Australia; Kids Research, Children’s Cancer Research Unit, The Children’s Hospital at Westmead, Westmead, NSW, Australia

## Abstract

Proteins are ultimately responsible for cellular phenotypes and are targeted by most anticancer drugs. However, beyond immunohistochemistry, proteins are not typically measured in precision oncology, meaning transcriptomics is used as a proxy. To determine how informative mRNA is for guiding personalised treatments, mRNA-protein correlations were analysed in three large pan-cancer datasets and made available in a web portal (https://procan.shinyapps.io/OnCorr/). *OnCorr* can be integrated into precision medicine programs to augment transcriptomics.

## Main text

Precision medicine programs in oncology almost universally incorporate genomic data, usually via targeted sequencing panels, or more rarely whole genome sequencing, to detect somatic variants that aid diagnosis or choice of therapy^1^. Some precision medicine initiatives also concurrently measure the transcriptome, with gene expression used to identify cancer subtypes or the upregulation of genes whose translated proteins correspond to known anti-cancer drug targets^2–4^. However, few precision medicine initiatives routinely incorporate proteomic data at scale^5,6^. This is a potential issue because most targeted anti-cancer agents that are recommended for personalised treatments directly inhibit a protein or pathway in a cancer cell^7^.

Due to the paucity of proteomic data generated in precision oncology, ultimately gene expression must be relied upon as a proxy for protein abundances – which typically remain unmeasured in a patient’s cancer. However, gene expression measurements are generally only moderately correlated with protein abundances, with correlations approximating 0.4-0.5^8–12^. Both technical and biological factors can influence mRNA-protein correlations, such as post-transcriptional regulation including protein degradation and differing rates of mRNA translation^13^. Therefore, having accurate knowledge of how well mRNA expression levels predict protein abundances for any given gene would enhance decision-making in precision medicine. Here we present a freely available precision oncology tool called *OnCorr* to interrogate mRNA-protein correlations across cancer types, available at https://procan.shinyapps.io/OnCorr/.

To assess pan-cancer mRNA-protein correlations, we first interrogated a multi-omic dataset of 949 human cell lines spanning 6,692 proteins across over 40 cancer types^8^. This dataset (called ProCan-DepMapSanger) includes proteomics from data-independent acquisition mass spectrometry, acquired in a standardised laboratory^14,15^ across several technical replicates per cell line. The transcriptomics are from RNA-sequencing data available at Cell Model Passports^16,17^. First, the median Spearman’s mRNA-protein correlation was calculated for each gene-protein pair in this dataset across 19 tissue types that comprised measurements in at least 10 cell lines (**Fig. 1a**). mRNA-protein correlations in each tissue type ranged from 0.31-0.45, with a cohort-wide median gene-wise mRNA-protein correlation of 0.42, across 6,205 genes with both mRNA and protein measurements (**Fig. 1b**).

**Figure 1.**
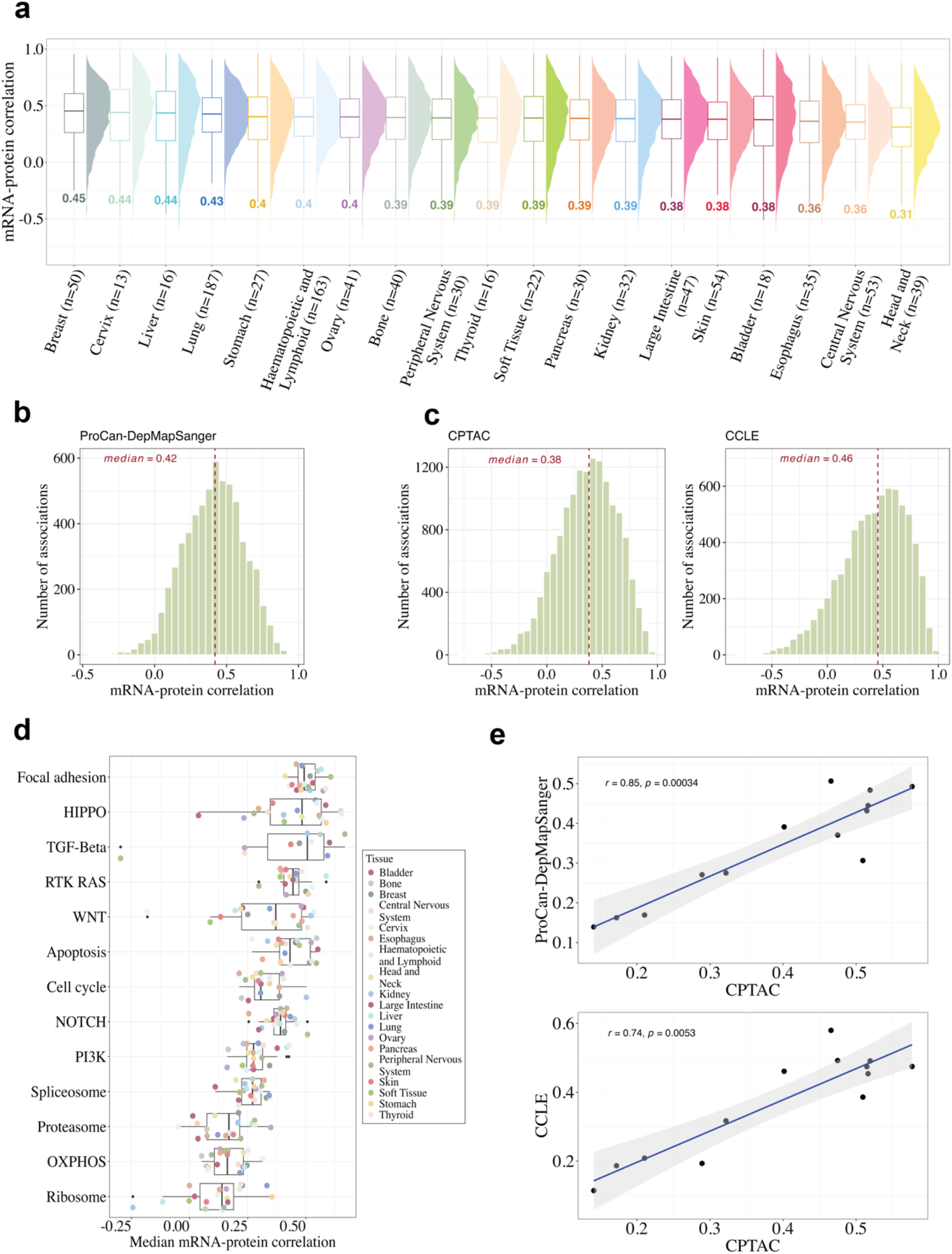
mRNA-protein correlations across tissues and cancer-related pathways. **a** Distribution of mRNA-protein correlations across tissues in the ProCan-DepMapSanger dataset. Only tissues with at least 10 cell lines are shown. **(b, c)** Histogram of mRNA-protein correlations across each cohort for **b** ProCan-DepMapSanger, **c** Clinical Proteomic Tumor Analysis Consortium (CPTAC; left) and Cancer Cell Line Encyclopedia (CCLE; right) datasets, with median indicated. **d** Boxplot showing median mRNA-protein correlations by tissue in the ProCan-DepMapSanger dataset for key biological and selected cancer-related pathways from Sanchez-Vega et al 23 and KEGG24, displayed similarly to data shown in Ghoshdastider et al 13. **e** Correlation of median mRNA-protein correlations per tissue for CPTAC with ProCan-DepMapSanger (left) and with CCLE (right), for pathways in **Fig. d** and **Supplementary Figure 1c**.

Next, mRNA-protein correlations from the ProCan-DepMapSanger cell line dataset were compared with two other publicly available pan-cancer studies with both transcriptome and proteome data. These include a series of cancer tissue cohorts from the Clinical Proteomic Tumor Analysis Consortium (CPTAC) (*n* = 1,030 cancer samples available from LinkedOmics^18^), and a pan-cancer cell line dataset from the Cancer Cell Lines Encylopedia (CCLE) (*n* = 375 cell lines)^9^. Both datasets show a similar range in mRNA-protein correlations across tissue types (0.32 to 0.49; **Supplementary Figure 1a** and **Supplementary Figure 1b**), with cohort-wide median correlations of 0.38 and 0.46 respectively (*n* = 14,465 and 10,437 genes; **Fig. 1c**). While cancer cell lines do not always capture the underlying biology of patient tissue^7^, they do have the advantage of not being impacted by issues relating to tumour purity, which could have a small effect on mRNA-protein correlation^13^. The cell line datasets selected for this work were all measured in a single study and, in the case of the ProCan-DepMapSanger dataset, on a single instrument platform, which allows for robust comparisons of correlations between tissue types. Increasing confidence in these datasets, both cancer cell line studies showed consistent mRNA-protein correlations across key biological pathways and selected cancer-related pathways as has been previously described in cancer tissues^13^ including those from CPTAC (Correlation of CPTAC with ProCan-DepMapSanger of *r* = 0.85 and with CCLE of *r* = 0.74; **Fig. 1d-e, Supplementary Figure 1c-d**).

mRNA-protein correlations were next interrogated across genes that often act as cancer drivers, using a set of 1,161 cancer genes from OncoKB^19^. This set combines genes from cancer sequencing panels, the COSMIC Cancer Gene Census^20^ and Vogelstein (2013)^21^. For robust analyses, mRNA-protein correlations were calculated only for the subset of these cancer driver genes in which the encoded protein was measured in at least ten samples in a minimum of fifteen tissue types (i.e., missing in fewer than five tissues; *n* = 261 genes). Using the ProCan-DepMapSanger dataset, hierarchical clustering revealed five clusters of driver genes with differing ranges of mRNA-protein correlations (**Fig. 2a**). Cancer drivers had slightly higher median correlations than genes not identified as cancer drivers (median *r* = 0.41 and 0.39, respectively; P < 0.0001 by unpaired Student’s t-test) (**Fig. 2b**). Among cancer drivers, one cluster of 50 genes had the highest mRNA-protein correlations across tissue types (Cluster 4; median *r* = 0.69) and is enriched for genes from pathways involved in chromatin binding and chromosome organisation (**Fig. 2c**). Another cluster of 12 genes had the poorest mRNA-protein correlations (Cluster 1; Median *r* = 0.16) and is comprised of genes from pathways involved in transcription factor and RNA polymerase II binding (**Fig. 2c**). Other clusters had genes with mRNA-protein correlations that varied considerably between tissue types. These patterns of mRNA-protein correlations among cancer driver genes could be replicated in the cancer tissue dataset from CPTAC (**Supplementary Figure 2a**) and the cancer cell line dataset from CCLE (**Supplementary Figure 2b**).

**Figure 2.**
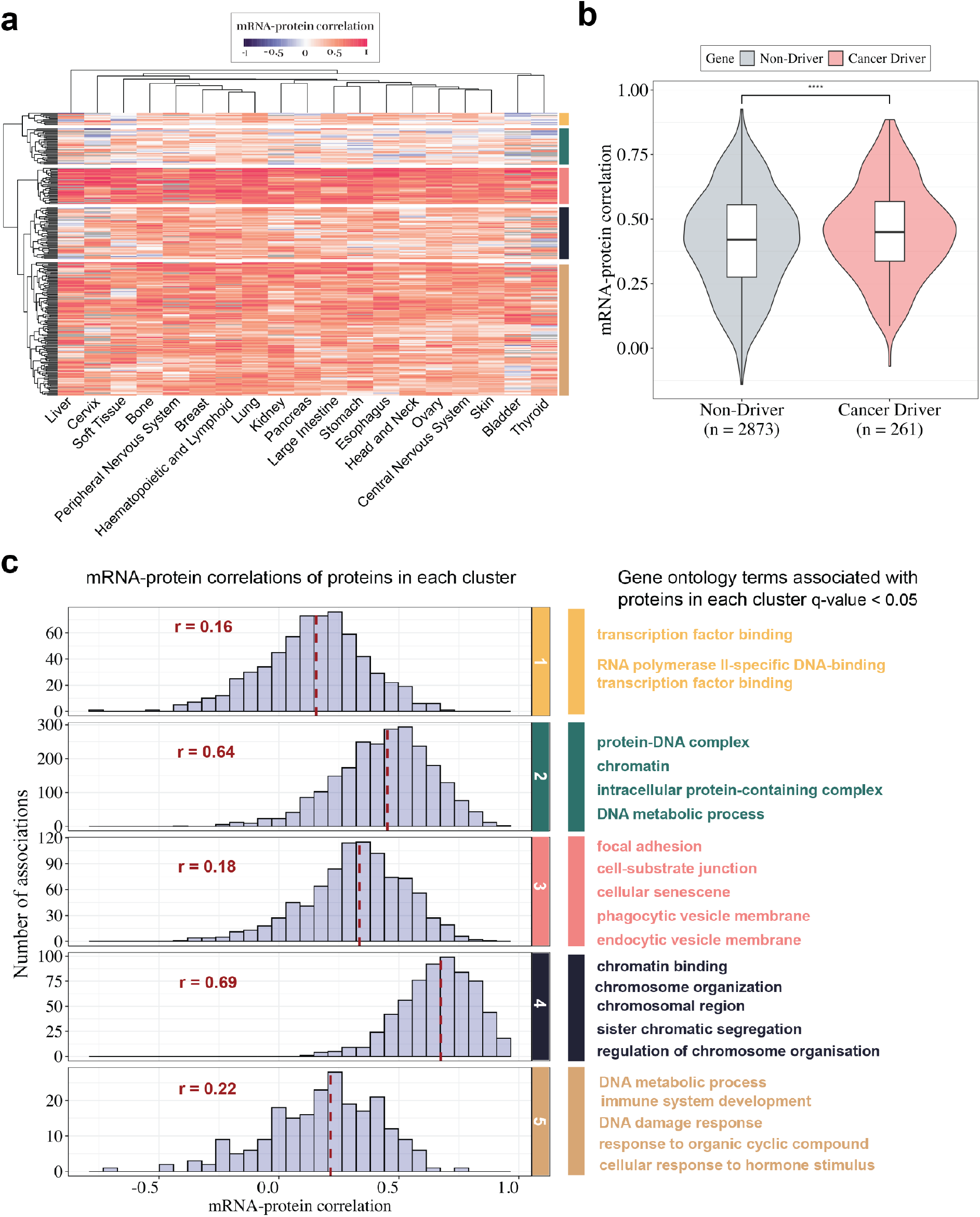
mRNA-protein correlations of cancer driver genes from OncoKB in the ProCan-DepMapSanger Dataset. **a** Heatmap of mRNA-protein correlations across tissue types for cancer driver genes from OncoKB^19^, with clusters identified by hierarchical clustering using Euclidean distance. **b** Violin plot showing mRNA-protein correlations of cancer driver and non-driver genes with median and standard deviation indicated. **** indicates P < 0.0001 by unpaired Student’s t-test. **c** Distribution of mRNA-protein correlations for genes in each cluster identified in **a**. Gene ontology terms with q-value < 0.05 are shown for each cluster. For all plots, only genes observed in at least ten samples in a minimum of fourteen tissues (i.e., missing in five or fewer tissues) are considered.

The extent of correlation between any given gene and the protein that it encodes can be driven by both technical and biological factors^12,13^. Upadhya et al (2022)^22^ developed a reproducibility rank to quantify the accuracy by which a protein can be measured by mass spectrometry. This ranking was shown to have a strong relationship with mRNA-protein correlation, suggesting that technical factors can influence the interpretation of such correlations. This association can also be observed in the ProCan-DepMapSanger (r = 0.49, **Fig. 3a**), CPTAC (r = 0.40, **Supplementary Figure 3a**) and CCLE (r = 0.29, **Supplementary Figure 3b**) datasets. Therefore, this reproducibility rank is a useful aid in the interpretation of mRNA-protein correlations, as an indicator of confidence in protein measurement accuracy across datasets and platforms.

**Figure 3.**
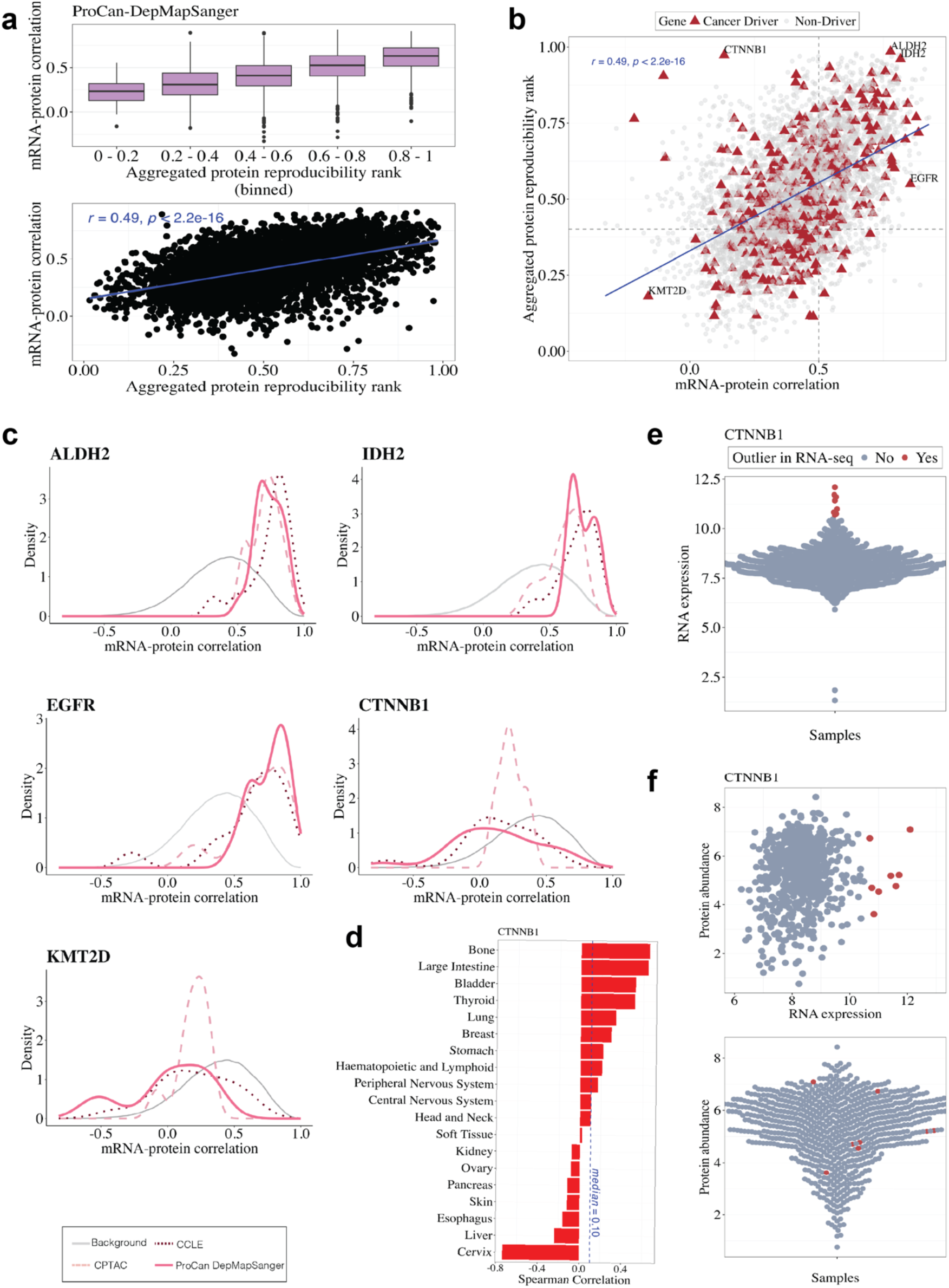
Integration of protein reproducibility ranks and gene-specific mRNA-protein correlations. **a** Association between aggregated protein reproducibility ranks from Upadhya et al 22 and mRNA-protein correlations in the ProCan-DepMapSanger dataset. Aggregated protein reproducibility ranks are binned (upper), with the Spearman’s correlation from unbinned data shown (lower). **b** Scatter plot of mRNA-protein correlation against aggregated protein reproducibility rank, with cancer drivers indicated by triangle data points. Labels indicate genes highlighted in the text and shown in **c. c** For each gene indicated in **b**, plots show the distribution of mRNA-protein correlations across tissues in the ProCan-DepMapSanger (pink solid line), Clinical Proteomic Tumor Analysis Consortium (CPTAC; light dashed line) and Cancer Cell Line Encyclopedia (CCLE; dark dashed line) datasets. Background (grey solid line) indicates the median mRNA-protein correlation from all genes in the ProCan-DepMapSanger dataset. Only tissues with data from a minimum of ten samples are included. **d** mRNA-protein correlation of CTNNB1 for each tissue type. **e** Distribution of CTNNB1 mRNA expression across samples from the ProCan-DepMapSanger dataset, with outliers (> 3 standard deviations above the mean) indicated. **f** Scatterplot showing mRNA expression and protein abundance for CTNNB1 (upper) and distribution of CTNNB1 protein abundance (lower) across samples in the ProCan-DepMapSanger dataset. Outliers indicated in **f** are those calculated from mRNA expression data in **e**.

To demonstrate the utility of mRNA-protein correlations for precision medicine, 474 cancer driver genes with robust measurements (i.e., observed with a minimum of 10 measurements from at least one tissue type) were interrogated in detail. Of these, 129 cancer driver genes had an mRNA-protein correlation above 0.5 and a good protein reproducibility rank (> 0.4; **Fig. 3b**). For these genes, which include ALDH2, IDH2 and EGFR (**Fig. 3c**), their transcriptome measurements should typically be considered a good predictor of protein abundance. In contrast, 30 cancer driver genes had mRNA-protein correlations below 0.25 while maintaining a good protein reproducibility rank (> 0.4; **Fig. 3b**). For these genes, which include CTNNB1 (**Fig. 3c**), transcriptome measurements will typically be a poor predictor of protein abundance. Proteins translated from such genes may be unreliable targets when identified from gene expression alone within a precision oncology pipeline. To explore this further with CTNNB1 as an example, a poor mRNA-protein correlation (median *r* = 0.10) is observed in most tissues (**Fig. 3d**). This suggests that in many tissues, outliers identified by RNA-sequencing (> 3 standard deviations above the mean; **Fig. 3e**) are unlikely to be highly expressed at the protein level (see outlier samples by RNA-seq in **Fig. 3f**). Finally, other genes such as KMT2D (**Fig. 3c**), have both low mRNA-protein correlations and poor reproducibility ranks. These genes have uncertain implications for precision oncology as a low mRNA-protein correlation may be a result of either biological factors such as posttranslational modifications or technical factors that influence protein measurement accuracy.

To make these mRNA-protein correlations accessible for precision medicine programs, the *OnCorr* web portal was built (https://procan.shinyapps.io/OnCorr/). This platform enables the interrogation of mRNA-protein correlations alongside reproducibility ranks within an easy-to-navigate user interface (**Fig. 4**). The tool integrates the ProCan-DepMapSanger, CPTAC and CCLE pan-cancer datasets, allowing the user to select their dataset of interest, as well as to filter and view results across all samples and by tissue type. This web portal is freely available and is designed to be interpretable by oncologists, researchers and data curators.

**Figure 4.**
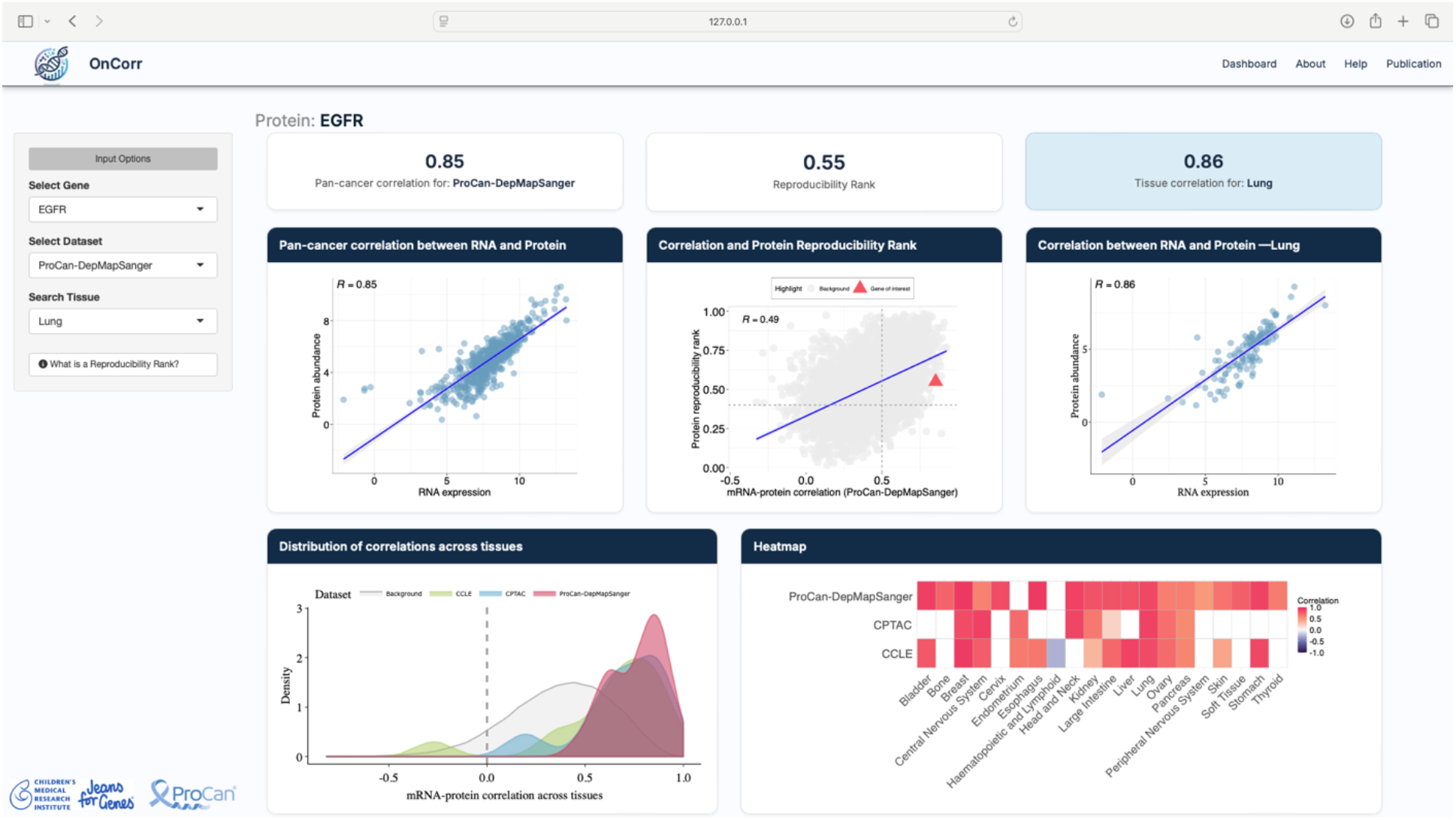
Example output from OnCorr web tool. A screenshot from the *OnCorr* web tool showing mRNA-protein correlations for EGFR, with tissue-specific data from lung.

This study and associated data interpretation have several limitations. For some genes, different mRNA-protein correlations are observed across datasets. This could be a result of differences in cell culture conditions, proteomic data acquisition methods or differences in the cancer types that make up each tissue annotation. Also, many of the datasets included in this study compare RNA-sequencing data acquired from a temporally or spatially different specimen than was used for the mass spectrometry analyses, leading to a false reduction in the measured mRNA-protein correlation. Finally, some genes, such as CDK4 (**Supplementary Figure 3c**), show patient outliers in mRNA expression (**Supplementary Figure 3d**) that remain highly expressed at the protein level (**Supplementary Figure 3e**) despite what would be suggested by the overall mRNA-protein correlation and reproducibility rank for these genes (CDK4: *r* = 0.14 correlation and 0.51 reproducibility rank). Therefore, inclusion of mRNA-protein correlations in precision oncology should be considered as one piece of information alongside several other metrics that, together, will drive the choice of targeted agents from a patient’s molecular profile.

In summary, *OnCorr* can be used in precision oncology initiatives to augment transcriptomic findings in the absence of proteomic data to inform a patient’s personalised treatment options. It can also be adopted for research to interrogate mRNA-protein correlations across three independent pan-cancer datasets. *OnCorr* is available at https://procan.shinyapps.io/OnCorr/.

## Methods

### Raw data sources

All data analysed in this study are publicly available from three independent datasets. Data from ProCan-DepMapSanger are available from Gonçalves et al ^8^. Data from the Clinical Proteomics Tumor Analysis Consortium (CPTAC) were downloaded from the LinkedOmicsKB database^18^. Data from the Cancer Cell Line Encyclopedia (CCLE) are available from Nusinow et al ^9^. Tissue type annotations were harmonised as in **Supplementary Table 1**.

### Correlation analysis

To calculate mRNA-protein correlations, a spearman rank correlation was performed for each mRNA-protein pair across all samples. Analyses were restricted to proteins with measurements in at least 10 cell lines for both RNA-seq and proteomics datasets to reduce any biases arising from proteins with measurements only in very few samples. For correlation analysis of mRNA-protein pairs within each tissue, cancers were grouped based on their tissue of origin and analysis was restricted to tissues that contained at least 10 cell lines in the relevant dataset.

### Functional enrichment

For biological pathways shown in **Fig. 1d** and **Supplementary Figure 1c**, pathways and their associated genes were retrieved from Sanchez-Vega et al ^23^ and KEGG pathways^24^, where *Homo sapiens* genes were retrieved from the msigdbr package (https://cran.r-project.org/web/packages/msigdbr/index.html). The median Spearman correlation rank was calculated for genes in each pathway per tissue type.

### Analysis of cancer driver genes

A list of 1,169 cancer driver genes was obtained from the OncoKB portal^19^. For heatmap and clustering analyses, genes were retained if they had mRNA-protein correlations in ≥15 tissue types (i.e., missing in fewer than 5 out of 19 tissues). The heatmap at **Fig. 2c** was plotted using the pheatmap package (https://cran.r-project.org/web/packages/pheatmap/index.html), which uses complete hierarchical clustering, and Euclidean distance as a similarity measure. The function *cutree* with the option k=5 was used to retrieve the driver genes in each cluster. Gene ontology analysis for each cluster was performed using clusterProfiler^25^. Only ontologies that had an FDR adjusted p-value < 0.05 were retained.

### Correlation analysis with reproducibility ranks

Aggregated protein-protein reproducibility ranks were retrieved from Upadhya et al ^22^. Ranks were binned for plotting and Spearman’s correlations were reported from unbinned data.

### Statistical analyses

All analyses were carried out in Python version 3.12.1, and R version 4.3.2, using the pandas library (Python), and dplyr, tidyverse and reshape2 (R). All plots were made using ggplot2 (https://cran.r-project.org/web/packages/ggplot2/index.html), and extensions ggdist (https://cran.r-project.org/web/packages/ggdist/index.html), ggpubr (https://cran.r-project.org/web/packages/ggpubr/index.html) and ggbeeswarm (https://cran.r-project.org/web/packages/ggbeeswarm/index.html).

## Data and code availability

The datasets analysed during the current study are available in Gonçalves et al ^8^, the LinkedOmicsKB database^18^ and Nusinow et al ^9^. The underlying code for this study is available in GitHub and can be accessed via www.github.com/CMRI-procan/OnCorr.

## Acknowledgements

ProCan is supported by the Australian Cancer Research Foundation, Cancer Institute New South Wales (NSW) (2017/TPG001, REG171150, 2021/CBG0002), NSW Ministry of Health (CMP-01), the University of Sydney, Cancer Council NSW (IG 18-01), Ian Potter Foundation, the Medical Research Future Fund (MRFF-PD), National Health and Medical Research Council (NHMRC) of Australia European Union grant (GNT1170739, a companion grant to support the ‘iPC-individualized Paediatric Cure’ [ref. 826121]), and National Breast Cancer Foundation (IIRS-18-164). Work at ProCan is done under the auspices of a Memorandum of Understanding between Children’s Medical Research Institute and the U.S. National Cancer Institute’s International Cancer Proteogenome Consortium (ICPC) that encourages cooperation among institutions and nations in proteogenomic cancer research in which datasets are made available to the public. R. C. P. and B. P. are supported by a Sydney Cancer Partners Translational Partners Fellowship with funding from a Cancer Institute NSW Capacity Building Grant (grant ID 2021/CBG0002). L. M. S. L. is funded by a CINSW Program Grant (no. 2021/TPG2112) and NHMRC Synergy Grant (APP2018642). This work was supported by NHMRC (GNT2000855, GNT1138536).

## Author contributions

R.C.P designed and directed the project. U. N. and R. C. P. analysed the data and wrote the manuscript. U.N. and O.L. built the *OnCorr* web tool. N. D. contributed statistical oversight of analyses. C.M., L. M. S. L., R. R. R., B. P. and R.C.P. interpreted the results and the implications for clinical implementation. All authors discussed the results and contributed to the final manuscript.

## Competing interests

The authors declare no competing interests

## Notes

https://procan.shinyapps.io/OnCorr/

### Competing Interest Statement

The authors have declared no competing interest.

