## Supplementary Material for "*OnCorr:* A pan-cancer mRNA-protein correlation tool for precision oncology"

### Supplementary Figures

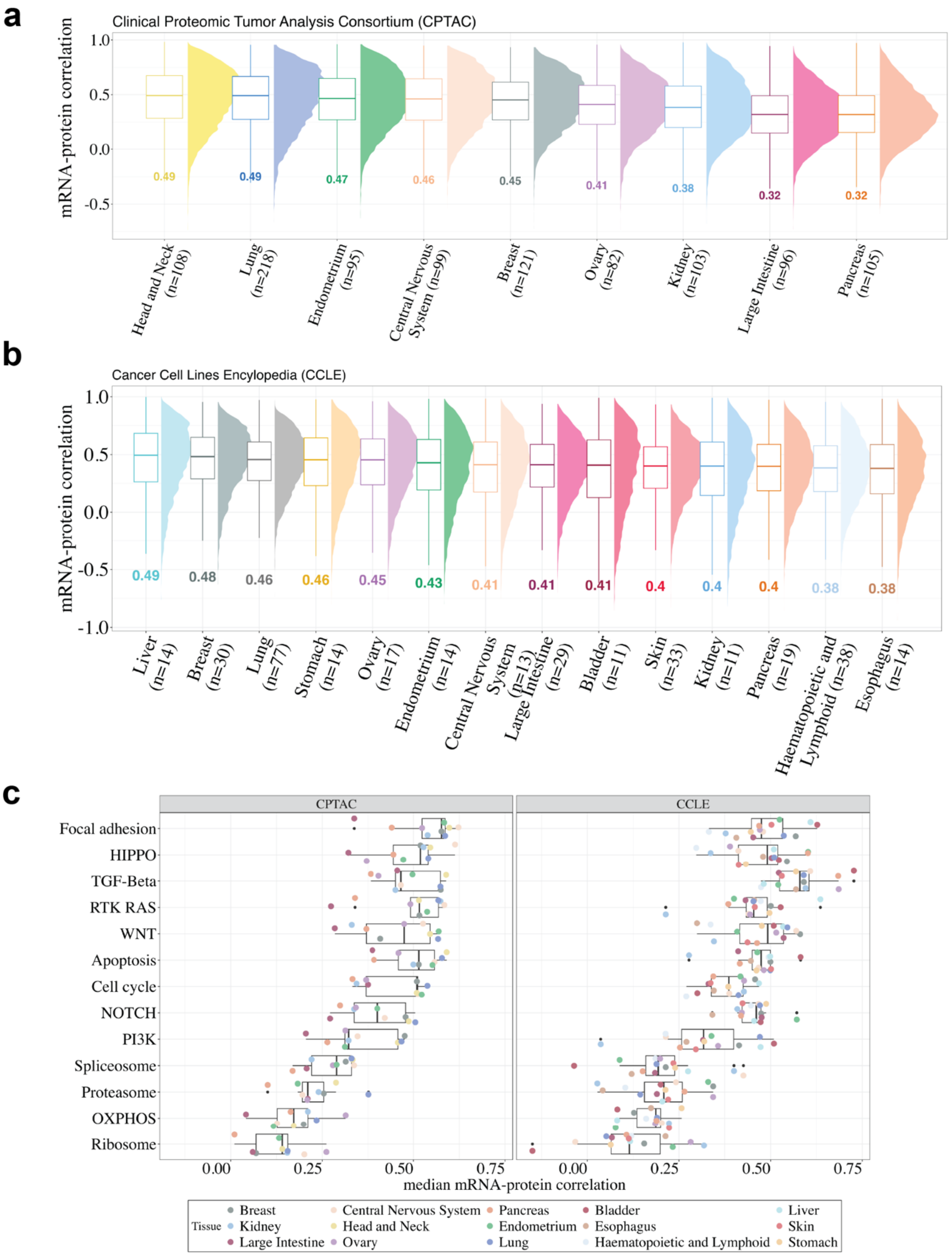

***Supplementary Figure 1. Overall mRNA-protein correlations across tissues and cancer-related pathways.*** **(a, b)** Distribution of mRNA-protein correlations across tissues in the **a** Clinical Proteomic Tumor Analysis Consortium (CPTAC) and **b** Cancer Cell Line Encyclopedia (CCLE) datasets. Only tissues with at least 10 cell lines are shown. **c** Boxplot showing median mRNA-protein correlations by tissue in the CPTAC (left) and CCLE (right) datasets for key biological and selected cancer-related pathways from Sanchez-Vega et al [23] and KEGG[24], displayed similarly to data shown in Ghoshdastider et al [13].

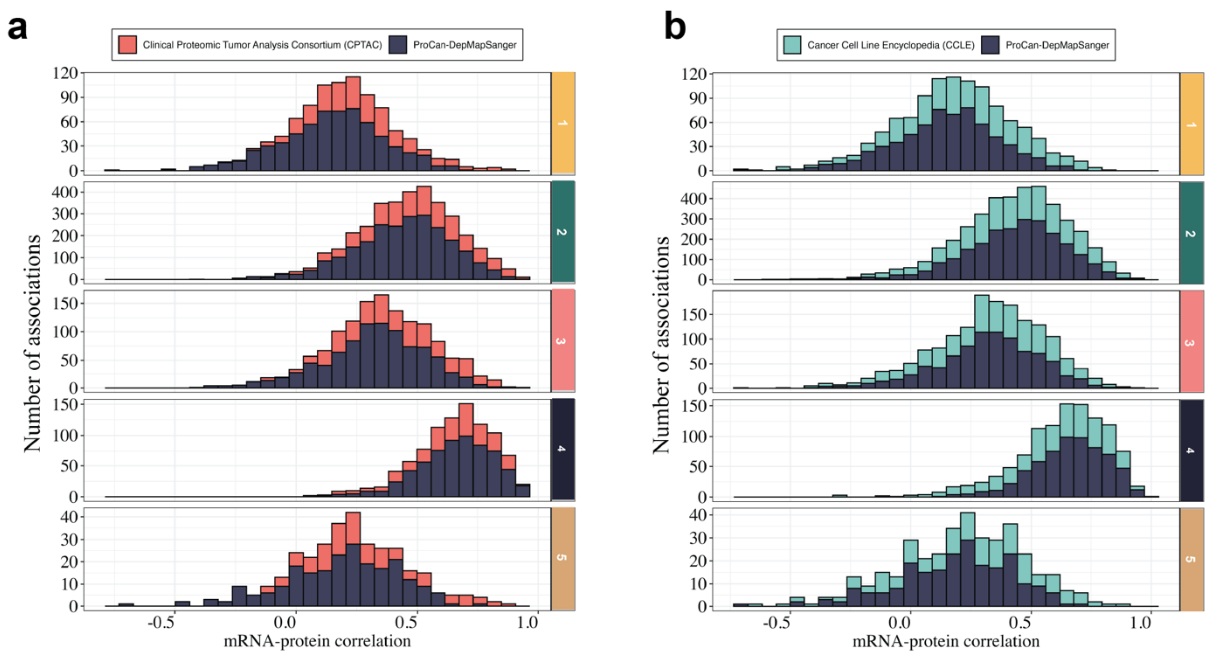

***Supplementary Figure 2. mRNA-protein correlations within clusters of cancer driver genes from the Clinical Proteomic Tumor Analysis Consortium (CPTAC) and Cancer Cell Line Encyclopedia (CCLE).*** Distribution of mRNA-protein correlations within tissue types for each cancer driver gene cluster identified in **Fig. 2a** for the **a** CPTAC and **b** CCLE datasets, each overlayed against the ProCan-DepMapSanger dataset. Only tissues with data from a minimum of ten samples are included.

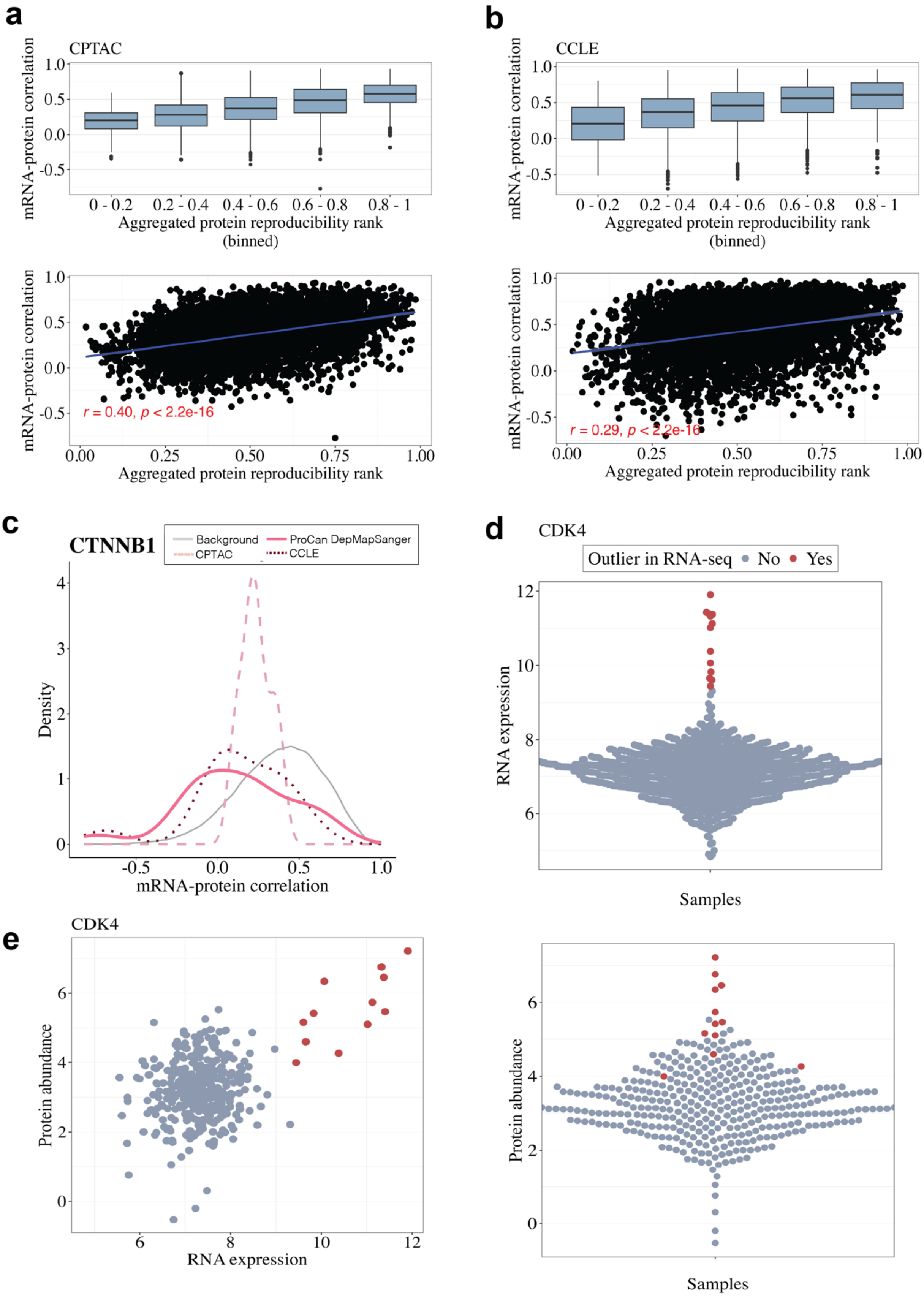

***Supplementary Figure 3. Protein reproducibility ranks across datasets and CDK4 associations.*** **(a, b)** Association between aggregated protein reproducibility ranks from Upadhya et al [22] and mRNA-protein correlations in the **a** Clinical Proteomic Tumor Analysis Consortium (CPTAC) and **b** Cancer Cell Line Encyclopedia (CCLE) datasets. Aggregated protein reproducibility ranks are binned (upper), with the Spearman's correlation from unbinned data shown (lower). **c** Distribution of mRNA-protein correlations across tissues for CDK4 in the ProCan-DepMapSanger (pink solid line), Clinical Proteomic Tumor Analysis Consortium (CPTAC; light dashed line) and Cancer Cell Line Encyclopedia (CCLE; dark dashed line) datasets. Background (grey solid line) indicates the median mRNA-protein correlation from all genes in the ProCan-DepMapSanger dataset. Only tissues with data from a minimum of ten samples are included. **d** Distribution of CDK4 mRNA expression across samples from the ProCan-DepMapSanger dataset, with outliers (> 3 standard deviations above the mean) indicated. **e** Scatterplot showing mRNA expression and protein abundance for CDK4 (left) and distribution of CDK4 protein abundance (right) across samples in the ProCan-DepMapSanger dataset. Outliers indicated in **e** are those calculated from mRNA expression data in **d**.

### Supplementary Tables

**Supplementary Table 1 –** Harmonisation of tissue type annotations across studies.

| Tissue Type Label in this study | Label in ProCan-DepMapSanger – Goncalves et al | Label in CPTAC (from LinkedOmics) | Label in CCLE – Nusinow et al |
|---|---|---|---|
| Adrenal Gland | Adrenal Gland | | adernal_cortex |
| Biliary Tract | Biliary Tract | | bile_duct |
| Bladder | Bladder | | urinary_tract |
| Bone | Bone | | bone |
| Breast | Breast | BRCA | breast |
| Central Nervous System | Central Nervous System | GBM | central_nervous_system |
| Cervix | Cervix | | cervix |
| Embryo | | | embryo |
| Endometrium | Endometrium | UCEC | uterus |
| Esophagus | Esophagus | | esophagus |
| Eye | | | eye |
| Fibroblast | | | fibroblast |
| Haematopoietic and Lymphoid | Haematopoietic and Lymphoid | | blood |
| Haematopoietic and Lymphoid | | | lymphocyte |
| Haematopoietic and Lymphoid | | | plasma_cell |
| Head and Neck | Head and Neck | HNSCC | upper_aerodigestive |
| Kidney | Kidney | CCRCC | kidney |
| Large Intestine | Large Intestine | COAD | colorectal |
| Liver | Liver | | liver |
| Lung | Lung | LUAD | |
| Lung | Lung | LSCC | lung |
| Ovary | Ovary | OV | ovary |
| Pancreas | Pancreas | PDAC | pancreas |
| Peripheral Nervous System | Peripheral Nervous System | | peripheral_nervous_system |
| Placenta | Placenta | | |
| Prostate | Prostate | | prostate |
| Skin | Skin | | epidermoid_carcinoma |
| Skin | Skin | | skin |
| Small Intestine | Small Intestine | | |
| Soft Tissue | Soft Tissue | | soft_tissue |
| Stomach | Stomach | | gastric |
| Testis | Testis | | |
| Thyroid | Thyroid | | thyriod |
| Unknown | | | unknown |
| Vulva | Vulva | | |
| | Other tissue | | |
